# A network-based approach for isolating the chronic inflammation gene signatures underlying complex diseases towards finding new treatment opportunities

**DOI:** 10.1101/2022.02.10.479987

**Authors:** Stephanie L Hickey, Alexander McKim, Christopher A Mancuso, Arjun Krishnan

## Abstract

Complex diseases are associated with a wide range of cellular, physiological, and clinical phenotypes. To advance our understanding of disease mechanisms and our ability to treat these diseases, it is critical to delineate the molecular basis and therapeutic avenues of specific disease phenotypes, especially those that are associated with multiple diseases. Inflammatory processes constitute one such prominent phenotype, being involved in a wide range of health problems including ischemic heart disease, stroke, cancer, diabetes mellitus, chronic kidney disease, non-alcoholic fatty liver disease, and autoimmune and neurodegenerative conditions. While hundreds of genes might play a role in the etiology of each of these diseases, isolating the genes involved in the specific phenotype (e.g., inflammation “component”) could help us understand the genes and pathways underlying this phenotype across diseases and predict potential drugs to target the phenotype. Here, we present a computational approach that integrates gene interaction networks, disease-/trait-gene associations, and drug-target information to accomplish this goal. We apply this approach to isolate gene signatures of complex diseases that correspond to chronic inflammation and prioritize drugs to reveal new therapeutic opportunities.

## 1 Introduction

Inflammation is an organism’s response to invasion by pathogens or to cellular damage caused by injury, and inflammatory responses fall into two categories: acute and chronic. During an acute inflammatory response, resident immune cells detect a pathogen or damaged cells in a tissue and release proinflammatory compounds, cytokines and chemokines, which recruit additional immune cells circulating in the blood to the site of injury or infection. These cells ultimately contain or neutralize the original offending stimulus, limiting further damage to the host, and clear dead cells and debris promoting tissue repair (Rock and Kono 2008). Systemic chronic inflammation (CI) occurs when inflammatory responses do not resolve, resulting in persistent, low-grade immune activation that causes collateral damage to the affected tissue over time (Furman et al. 2019).

Complex disorders like cardiovascular diseases, diabetes, cancer, and Alzheimer’s disease are among the leading causes of death and disability among adults over 50 years of age, and all are associated with underlying systemic inflammation (Furman et al. 2019; Vos et al. 2020). Patients with systemic inflammation caused by autoimmune disorders are more likely to have another CI disorder like cardiovascular disease, type 2 diabetes mellitus, and certain types of cancer (Dregan et al. 2014; Armstrong, Harskamp, and Armstrong 2013; Yashiro 2014). Further, treating one chronic-inflammatory disease can reduce the risk of contracting another, suggesting a common underlying pathway (Fullerton and Gilroy 2016). For example, treating rheumatoid arthritis with tumor necrosis factor (TNF) antagonists lowers the incidence of Alzheimer’s disease and type II diabetes (Chou et al. 2016; Antohe et al. 2012). Quantifying the similarity between chronic inflammation signatures across complex diseases could increase our understanding of these relationships.

Common treatments for systemic inflammation, including non-steroidal anti-inflammatory drugs (NSAIDs), corticosteroids, and biologics like tumor necrosis factor (TNF) antagonists, can cause adverse effects when used long term. For instance, patients treated with corticosteroids or TNF antagonists have increased risk of infection (M. Shah et al. 2013; Rosenblum and Amital 2011; Murdaca et al. 2015), and corticosteroid use increases both the risk of fracture (Kanis et al. 2004; Mitra 2011) and the risk of developing type II diabetes (Blackburn, Hux, and Mamdani 2002). NSAIDs present a unique set of side effects, particularly in elderly patients, including gastrointestinal problems ranging from indigestion to gastric bleeding, and kidney damage (Marie R Griffin 1998; Marcum and Hanlon 2010; M. R. Griffin, Yared, and Ray 2000). Therefore, the search for better treatment options for CI is ongoing.

Here, we present a computational approach that uses a network-based strategy to isolate CI gene “signatures” specific to each disorder. We quantify the similarity between CI signatures across diseases and predicted and prioritized potential new treatment opportunities specifically targeting the ‘inflammation’ component of complex diseases. Importantly, our strategy can be used to pinpoint the gene signature of any phenotype underlying a complex disease and identify drugs to treat that specific phenotype.

## 2 Methods

### 2.1 Disease selection and disease-associated seed genes

#### 2.1.1 Complex and autoimmune diseases

We searched the literature (Furman et al. 2019; Dregan et al. 2014; Armstrong, Harskamp, and Armstrong 2013; Yashiro 2014; Chou et al. 2016) and selected 17 complex diseases associated with chronic inflammation (CI) and 9 autoimmune diseases. Some of these diseases are quite broad (i.e “Malignant neoplasm of lung”). To add more narrowly defined diseases to our list, we used the Human Disease Ontology (Schriml et al. 2019) to identify child terms of the diseases. For each original disease and child disease, we used curated annotations from DisGeNET to find disease-associated genes. To ensure that our disease gene sets were largely non-overlapping, we only retained one representative from every group of disease gene sets with ≥ 0.6 overlap (|*A* ∩ *B*|/ min (|*A*|, |*B*|)). This resulted in 10 autoimmune diseases and 37 complex diseases (**Table S1**).

#### 2.1.2 Non-disease traits

∼100 non-disease-traits that are unlikely to be related to SNPs associated with CI (i.e., handedness, coffee intake, and average household income) were hand selected from the list of traits with GWAS results from the UK Biobank (Sudlow et al. 2015) to be used as negative controls. Based on GWAS summary statistics from the Neale group (Abbot et al. n.d.), we used Pascal (Lamparter et al. 2016) (upstream and downstream windows of 50 KB with the sum-of-chi-squared statistics method; only autosomal variants) to associate genes with the non-disease traits. Genes with *p* < 0.001 were included as seed genes for that trait.

### 2.2 GenePlexus

To predict new genes associated with a set of input seed genes, we used GenePlexus, a tool that builds an L2-regularized logistic regression model using features from a gene interaction network (Liu et al. 2020). As input features, we used the adjacency matrices from STRING, STRING with only experimentally derived edges (STRING-EXP) (Szklarczyk et al. 2017), BioGRID (Stark et al. 2006), and ConsensusPathDB (Kamburov et al. 2013). For predicting disease genes, positive examples were disease/trait seed genes and negative example genes were generated by: (i) finding the union of all genes annotated to all diseases in DisGeNET(Piñero et al. 2020), (ii) removing genes annotated to the given seed genes, and (iii) removing genes annotated to any disease in the collection that significantly overlapped with the given seed genes (*p* < 0.05 based on the one-sided Fisher’s exact test) (Liu et al. 2020). We tested the performance of the above features for predicting new genes associated with our diseases and traits of interest using three-fold cross validation and only included diseases in subsequent analyses if the diseases/traits had ≥ 15 associated genes and median *log*_2_(*auPRC*⁄*prior*) ≥ 1. (i.e. the area under the precision-recall curve ‘auPRC’ is at least twice as much as expected by random chance ‘prior’(Liu et al. 2020)). See **Table S1**.

### 2.3 Identifying clusters of interacting genes within a disease-specific network

One list of disease-associated genes was formed for each of the four biological networks used as features in GenePlexus. Specifically, we added genes with a GenePlexus prediction probability of ≥ 0.80 on the network of interest to the original disease or trait seed gene list to create our final set of associated genes for each disease or trait for that network. We formed disease/trait-specific networks by subsetting a particular network to include only the disease/trait associated genes and any edges connecting those genes based on direct interactions (**Fig. 1A**). We tested five prediction network/ cluster network combinations: Genes predicted on each of the four networks were clustered on the same network, genes predicted on STRING were clustered on STRING-EXP. We then used the Leiden algorithm (Traag, Waltman, and van Eck 2019) to partition the disease/trait-specific networks into clusters. Specifically, we used the *leiden_find_partition* function from the leidenbase R package (v 0.1.3) (https://github.com/cole-trapnell-lab/leidenbase with 100 iterations and ModularityVertexPartition as the partition type. We retained clusters containing ≥ 5 genes.

**Figure 1:**
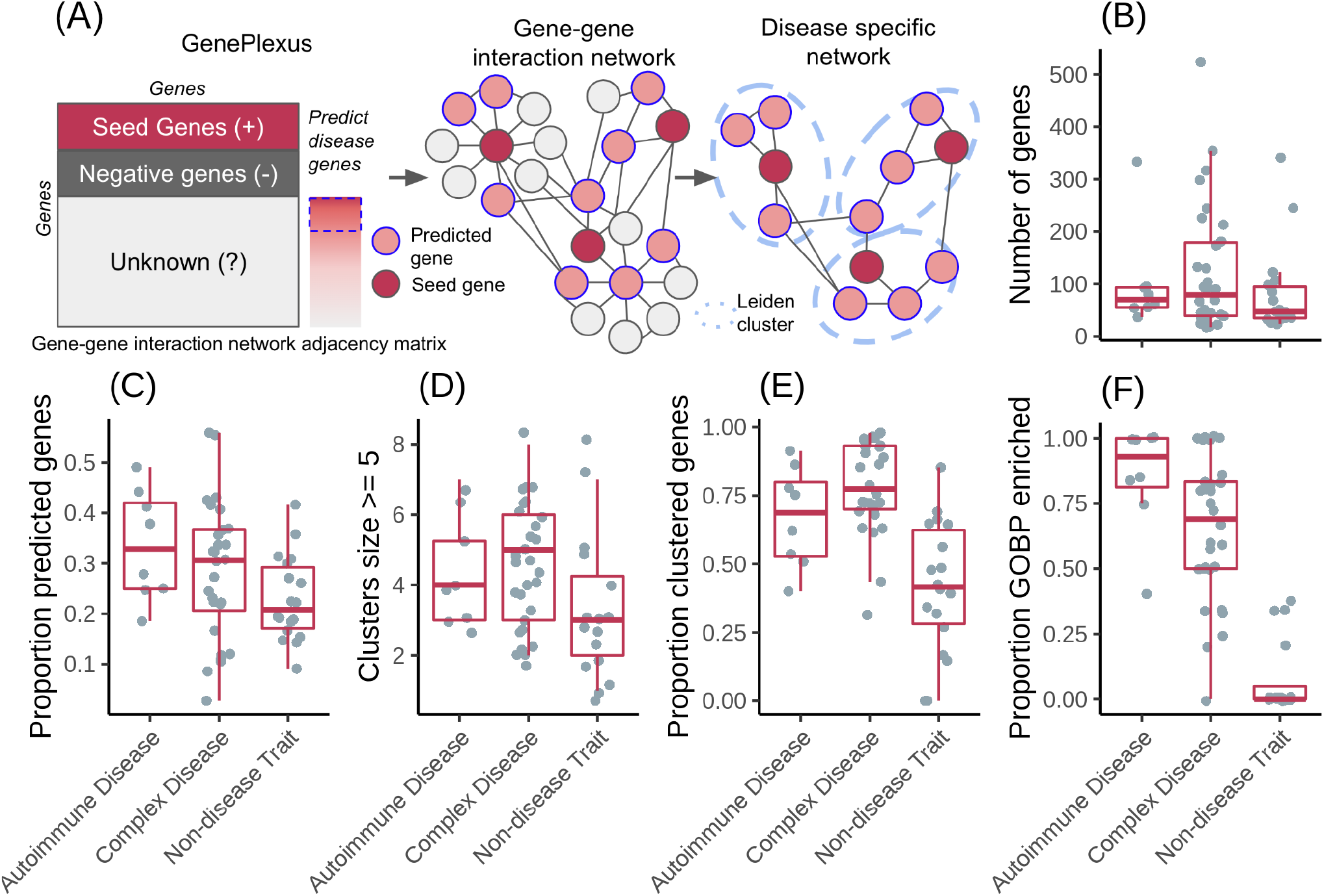
**(A)** Schematic outline of the expansion of disease-/trait-associated gene lists with GenePlexus and the clustering of trait specific networks. **(B)** Number of genes per disease/trait. **(C)** Proportion of the genes per disease/trait that were predicted by GenePlexus. **(D)** Number of clusters per disease/trait containing at least 5 genes. **(E)** Proportion of total genes assigned to a cluster containing at least 5 genes. **(F)** Proportion of clusters per disease/trait enriched with genes from at least one GO biological process.

### 2.4 Cluster GOBP enrichment analysis

We used the R package topGO with the “weight01” algorithm and Fisher testing (Alexa and Rahnenfuhrer 2022) (v 2.44.0) to find enrichment of genes annotated to GO biological processes (min size = 5, max size = 100) among disease gene clusters. The annotations were taken from the Genome wide annotation for Human bioconductor annotation package, org.Hs.eg.db (Carlson 2019) (v 3.13.0). The background gene set included all human genes present in the network of interest.

### 2.5 Isolating CI-associated disease clusters

#### 2.5.1 Defining CI-associated genes

We tested several different sets of chronic inflammation associated genes for this study including the GO biological process (GOBP) terms GO:0002544 (“chronic inflammatory response”) and GO:0006954 (“inflammatory response”). These were collected from the Genome wide annotation for Human bioconductor annotation package, org.Hs.eg.db (Carlson 2019) (v 3.13.0) with and without propagation of gene-term relationships from the descendent terms (org.Hs.egGO2ALLEGS and org.Hs.egGO2EG, respectively). GO:0006954 was also filtered to retain gene-term relationships inferred from experiments (evidence codes EXP, IDA, IPI, IMP, IGI, IEP, HTP, HDA, HMP, HGI, and HEP). As GO:0002544 without propagation contained < 15 genes, this list was ultimately not included in the study. We also identified genes associated with chronic inflammation using Geneshot which, given the search term “chronic inflammation”, searches Pubmed using manually collected GeneRif gene-term associations to return a ranked list containing genes that have been previously published in association with the search term (Lachmann et al. 2019). We tested both the entire Geneshot generated list, and the subset of genes with > 10 associated publications. As with the disease genes, we predicted additional chronic-inflammation-associated genes using GenePlexus with features from each adjacency matrix of interest. Negative examples for GenePlexus were derived from non-overlapping GOBP terms. We added genes with a prediction probability of ≥ 0.80 to the seed gene list to create our final sets of CI-associated genes.

#### 2.5.2 Creating random traits

After running GenePlexus to predict new genes for each trait, the gene lists for each trait were used to generate 5,000 random gene lists that have matching node degree distributions to the original traits. That is, a random gene list was generated for a given trait by replacing each of its genes in the network of interest with a (randomly chosen) gene that has the same node degree, or a gene that has a close node degree if there are a small number of genes with the exact node degree. We clustered the random traits as described in section 2.3. Only clusters with ≥ 5 genes were included. Real traits with no corresponding permuted traits with clusters containing ≥ 5 genes were excluded from the analysis.

#### 2.5.3 Finding CI-gene enriched disease clusters

For each prediction-network/cluster-network pair and each CI gene list expanded on the prediction network of interest, for each disease and random trait cluster containing ≥ 5 genes, we calculated an enrichment score, 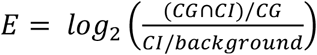 where CG are the genes in a disease cluster, CI are the CI genes, and background are all of the genes present in the clustering network. For each real disease or trait cluster, we used the matching random trait clusters to calculate a p-value, 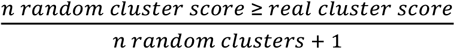. We corrected for multiple comparisons across clusters within a disease using the Benjamini-Hochberg procedure. Clusters with an *E* > 0 and *FDR* < 0.05 were considered chronic-inflammation-associated disease clusters and were deemed to represent the ‘CI signature’ of the disease.

### 2.6 Comparing CI-signatures across diseases

For CI-enriched clusters identified using ConsensusPathDB and the high-confidence Geneshot CI genes, we used the SAveRUNNER R package to quantify the similarity between each pair of CI-enriched clusters using ConsensusPathDB as the base network (Fiscon and Paci 2021). For each pair, SAveRUNNER computes the average shortest path between each gene in *cluster A* and the closest gene in *cluster B* and uses this value to calculate an adjusted similarity score. Then, a p-value is estimated based on a null distribution of adjusted similarity scores between randomly generated clusters with the same node degree distributions as clusters *A* and *B*. Because the similarity scores and p-values are not symmetric, i.e., *A* → *B* ≠ *B* → *A*, we used Stouffer’s method to combine p-values for the same pair of clusters and averaged the adjusted similarities. We then used the Leiden algorithm as described in section 2.3 to group related clusters. For each group, we took the union of the genes belonging to the resident CI-enriched clusters. Using genes unique to each group, with all the ConsensusPathDB genes as background, we used TopGO as in section 2.4 to identify enriched GOBPs.

### 2.7 Predicting novel treatment opportunities

#### 2.7.1 Identifying expert-curated drug-target associations

The known drug-gene interactions used in this study are the subset of the interactions present in the DrugCentral database (Avram et al. 2021) that are also among the expert curated interactions in the Drug-Gene Interaction database (DGIdb) (Freshour et al. 2021). Specifically, we used the DGIdb API to retrieve only drug-gene interactions that were marked “*Expert curated*” (based on the source trust levels endpoint). Intersecting these interactions with those in DrugCentral (through a list of drug synonyms from DrugCentral) resulted in the final list of expert-curated drug-gene pairs.

#### 2.7.2 Treatment prediction and scoring

We predicted treatment opportunities for the inflammatory component of complex diseases by using the SAveRUNNER R package (Fiscon and Paci 2021). SAveRUNNER builds a bipartite drug-disease network through utilizing the previously determined expert-curated drug targets, the CI-associated cluster disease genes, and the ConsensusPathDB network as a human interactome. Network similarity scores returned by SAveRUNNER represent the proximity between disease and drug modules, where a high similarity score means that the disease and drug modules have high proximity in ConsensusPathDB. SAveRUNNER calculates a p-value where a significant value represents the disease genes and drug targets are nearby in the network more than expected by chance (based on an empirical null distribution; see Section 2.6). Using the list of final predicted associations after normalization of network similarity, the p-values were corrected for multiple comparisons within each disease using the Benjamini-Hochberg procedure. Drugs were associated to diseases based on the disease cluster with the lowest FDR. Predicted treatments are disease-drug pairs with *FDR* < 0.01.

#### 2.7.3 Evaluating SAveRUNNER prediction performance

We calculated *log*_2_(*auPRC*⁄*prior*) by ranking disease-drug pairs by −*log*_10_(*SAveRUNNER FDR*) and using either previously indicated drug-disease pairs (both approved and off-label) or drug-disease pairs tested in a clinical trial as positive labels. Approved and off-label drug-disease pairs were collected from DrugCental (Avram et al. 2021). Only drugs with expert curated target genes were included (see section *2.6.1*). The Unified Medical Language System (UMLS) Concept Unique Identifiers (CUI) were limited to diseases (T047) and neoplastic processes (T191), and our diseases were matched to diseases in DrugCentral using UMLS CUI ids. Drug-disease pairs tested in a clinical trial were collected from the database for Aggregate Analysis of Clinical Trials (AACT) (“AACT Database | Clinical Trials Transformation Initiative” n.d.). AACT reports the Medical Subject Headings (MeSH) vocabulary names for diseases. We used disease vocabulary mapping provided by DisGeNET to translate UMLS CUI ids for our diseases to MeSH vocabulary names, further restricted to only those that were present in AACT. We filtered AACT for trials with “Active, not recruiting”, “Enrolling by invitation”, “Recruiting”, or “Completed” status.

## 3 Results

### 3.1 Expanding lists of disease-related genes and identifying disease-specific gene subnetworks

Our first goal was to establish a comprehensive list of genes associated with the complex diseases of interest and resolve the genes linked to each disease into subsets of tightly connected genes in an underlying molecular network.

Towards this goal, we selected 37 complex diseases associated with underlying systemic inflammation (see *Methods*). To ensure that we correctly isolate chronic inflammation (CI) signatures, we devised a set of positive and negative controls. We selected 10 autoimmune disorders as positive controls because autoimmune disorders are characterized by CI and should have an easily identifiable CI gene signature. For negative controls, we selected ∼100 traits from UK Biobank (Sudlow et al. 2015) that are unlikely to be associated with CI (i.e., Right handedness, filtered coffee intake, and distance between home and workplace). **Table S2** contains the full list of diseases and traits used in this analysis along with their original associated genes.

Our first goal was to establish a comprehensive list of genes associated with the complex diseases of interest and resolve the genes linked to each disease into subsets of tightly connected genes in an underlying molecular network. Towards this goal, we selected 37 complex diseases associated with underlying systemic inflammation (see *Methods*). To ensure that we correctly isolate chronic inflammation (CI) signatures, we devised a set of positive and negative controls. We selected 10 autoimmune disorders as positive controls because autoimmune disorders are characterized by CI and should have an easily identifiable CI gene signature. For negative controls, we selected ∼100 traits from UK Biobank (Sudlow et al. 2015) that are unlikely to be associated with CI (i.e., Right handedness, filtered coffee intake, and distance between home and workplace). **Table S2** contains the full list of diseases and traits used in this analysis along with their original associated genes.

Genes predicted by the GenePlexus model with a probability ≥ 0.80 were added to the seed genes to create an expanded list of disease-/trait-associated genes (**Fig. 1B**). The proportion of genes predicted by GenePlexus for the non-disease traits is lower than those for the autoimmune and complex diseases (**Fig. 1C**). This observation indicates that genes associated with a specific autoimmune/complex disease tend to have more similar network neighborhoods than genes associated with non-disease traits. All disease-associated genes after GenePlexus prediction are listed in **Table S3**.

Next, for each disease/trait, we clustered the expanded lists of genes based on their interaction in ConsensusPathDB. (**Fig.1A** and **D**; **Table S3**). The complex diseases had the highest proportion of genes grouped into clusters of ≥ 5 genes, followed by autoimmune diseases and non-disease traits (**Fig. 1E**). To assess whether clusters are biologically meaningful, we performed an enrichment analysis between every cluster and hundreds of GO Biological Process (GOBP) gene sets. We theorize that significant enrichment of a cluster with a GOBP means the genes in the cluster likely function together to carry out a specific cellular process or pathway. For autoimmune and complex diseases, the median proportion of GOBP enriched clusters are > 0.75 and > 0.60, respectively, suggesting most clusters are biologically relevant (**Fig. 1F**). In contrast, most clusters in non-disease traits are not enriched for a GOBP (**Fig. 1F**).

### 3.2 Isolating CI-enriched disease clusters

Clusters of related, disease-associated genes on functional gene interaction networks are likely to correspond to the pathways and biological processes disrupted during disease progression. For complex disorders, multiple pathways are likely to be affected. Our next goal was to identify which cluster(s) within a set of disease-associated genes corresponds to the CI component of the disease. For this analysis, similar to the expansion of disease-/trait-associated genes, we used GenePlexus to take the set of 133 human genes annotated to “chronic inflammation” by Geneshot via more than 10 publications (Lachmann et al. 2019) and expanded it to a list to high-confidence 235 CI genes (**Table S4**). We then scored the enrichment of CI genes in each disease cluster and performed a permutation test using 5,000 random gene sets for each disease to determine the significance of the enrichment score (see *Methods* section 2.5, **Table S5**). As expected, we were able to identify clusters enriched for CI genes in all the autoimmune disorders surveyed (9/9), while finding no CI-enriched clusters among the non-disease traits (**Fig. 2A**). We identified at least one CI-enriched cluster in 18 of 30 of the complex diseases (**Fig. 2A**). Twelve out of the 27 diseases with at least one CI-enriched cluster had two or more CI-enriched clusters, and the median proportion of CI-enriched clusters out of the total clusters is higher for autoimmune diseases than complex diseases (**Fig. 2B**).

**Figure 2:**
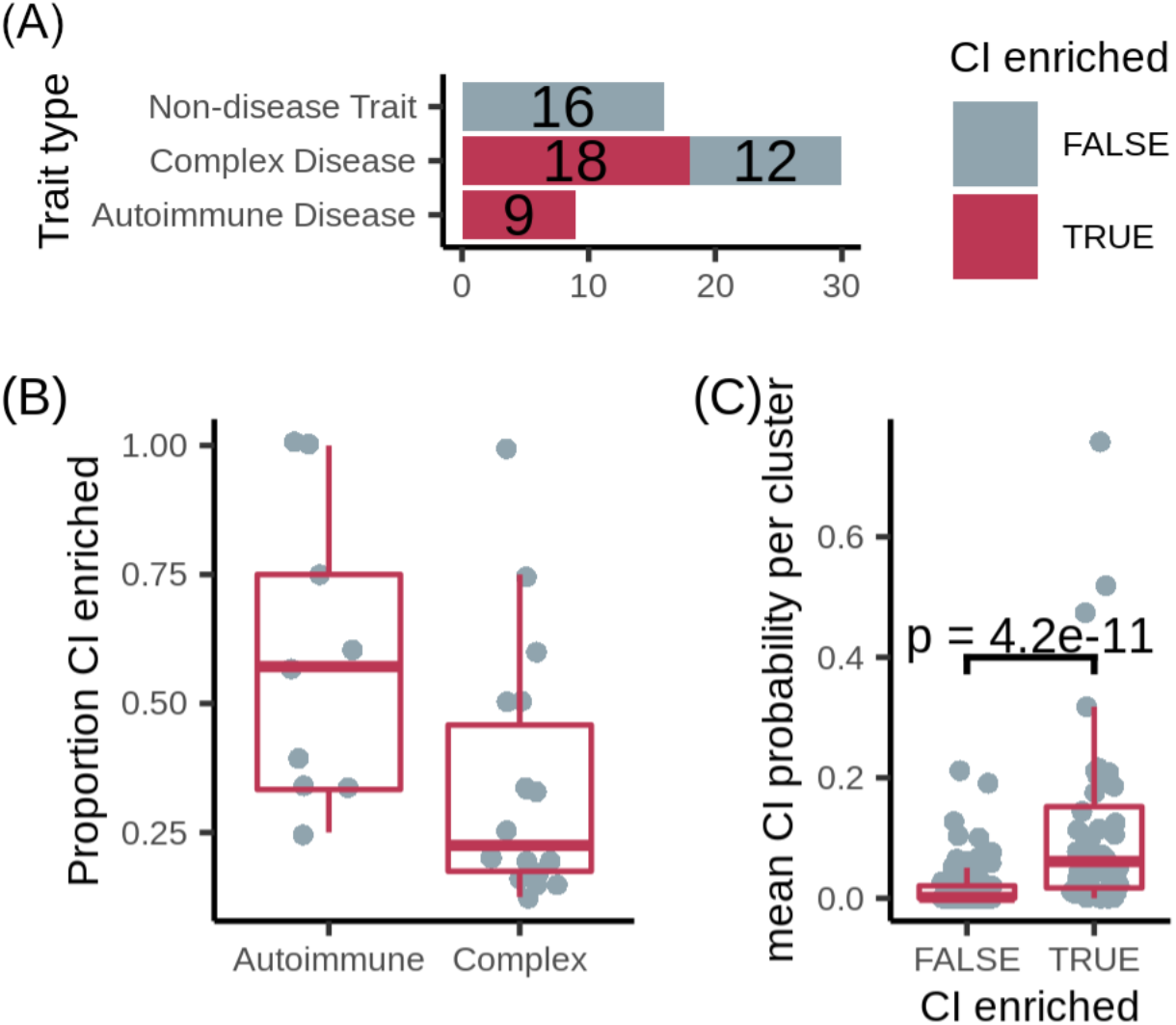
**(A)** Number of diseases/traits with at least one cluster overlapping the expanded chronic inflammation (CI) geneset (dark pink), out of the total number of diseases/traits. **(B)** The proportion of CI-enriched disease clusters among all disease clusters per disease. **(C)** Mean probability that genes with no known relationship with chronic inflammation residing in a CI-enriched cluster or non-CI-enriched cluster are associated with CI. P-value calculated using the Wilcoxon signed-rank test.

We hypothesized that, through guilt-by-association, even the genes with no known relationship with chronic inflammation residing in a CI-enriched cluster should have a higher probability of being CI-associated than those in non-CI-enriched clusters. To test this hypothesis, we used GenePlexus with features from ConsensusPathDB to calculate the probability that every gene is associated with high-confidence Geneshot CI gene set. Then, focusing on the genes in disease clusters that were not present in the high-confidence Geneshot CI gene set, we found that the mean CI probability of these genes in CI-enriched clusters is significantly higher for CI-enriched clusters than non-enriched clusters (**Fig. 2C**). This observation suggests that the CI-enriched clusters as a whole, and not just the genes in the high-confidence Geneshot CI gene set residing within them, are CI-associated.

To test the robustness of this method for identifying CI enriched clusters, we performed disease-gene expansion and clustering using three other biological interaction networks of varying sizes and edge densities — STRING, STRING with only experimentally derived edges (STRING-EXP) (Szklarczyk et al. 2017), and BioGRID (Stark et al. 2006) — and we overlapped the resulting disease subclusters with the 235 genes from the high-confidence Geneshot list as well as four additional lists of chronic inflammation associated genes expanded on the network of interest, including the full set of Geneshot genes and three sets of genes derived from the “chronic inflammatory response” and “inflammatory response” GOBP terms (see *Methods* section 2.5, **Tables S3-S5, Fig. S1**). Using ConsensusPathDB as the base network for predicting new disease genes and generating disease-specific subnetworks and overlapping these subnetworks with genes associated with the high-confidence Geneshot chronic inflammation gene set, resulted in the highest proportion of autoimmune diseases and lowest proportion of non-disease traits with at least one CI-enriched cluster of any base-network/CI-gene set combination. Nevertheless, while the number of diseases with at least one CI-enriched cluster varied with different combinations of prediction network, cluster network, and inflammation gene set, in every case (**Table S6**), the proportion of autoimmune diseases with at least one CI-enriched cluster was higher than that for non-disease traits (**Fig. S2**). Moreover, the mean CI probability of genes in CI-enriched clusters is significantly higher for CI enriched clusters than non-enriched clusters in 24 out of 25 cases (**Fig. S3-S7**), suggesting that our method is robust to changes in base-network and inflammation gene set.

### 3.3 Comparing CI gene signatures across diseases

To determine if related diseases have similar chronic inflammation signatures, we used a network-based approach to quantify the similarity between each pair of CI-enriched disease clusters across diseases and grouped similar clusters together using the Leiden algorithm (**Table S7**) (Traag, Waltman, and van Eck 2019; Fiscon et al. 2021). Several diseases have more than one CI-enriched cluster and none of these diseases have clusters belonging only to one group (**Fig. 3A**). Moreover, diseases belonging to the same broad category — i.e., autoimmune, cancer, or cardiovascular disease — do not have a larger proportion of clusters belonging to a particular group than expected by chance (one-sided Fisher’s exact test, **Fig. 3A**). This suggests that one disease can harbor more than one type of chronic-inflammation signature, and that the same signatures can be found in very different diseases. For example, rheumatoid arthritis, myocardial ischemia, atherosclerosis and chronic obstructive airway disease all have CI-enriched clusters belonging to each of the three signature groups.

**Figure 3:**
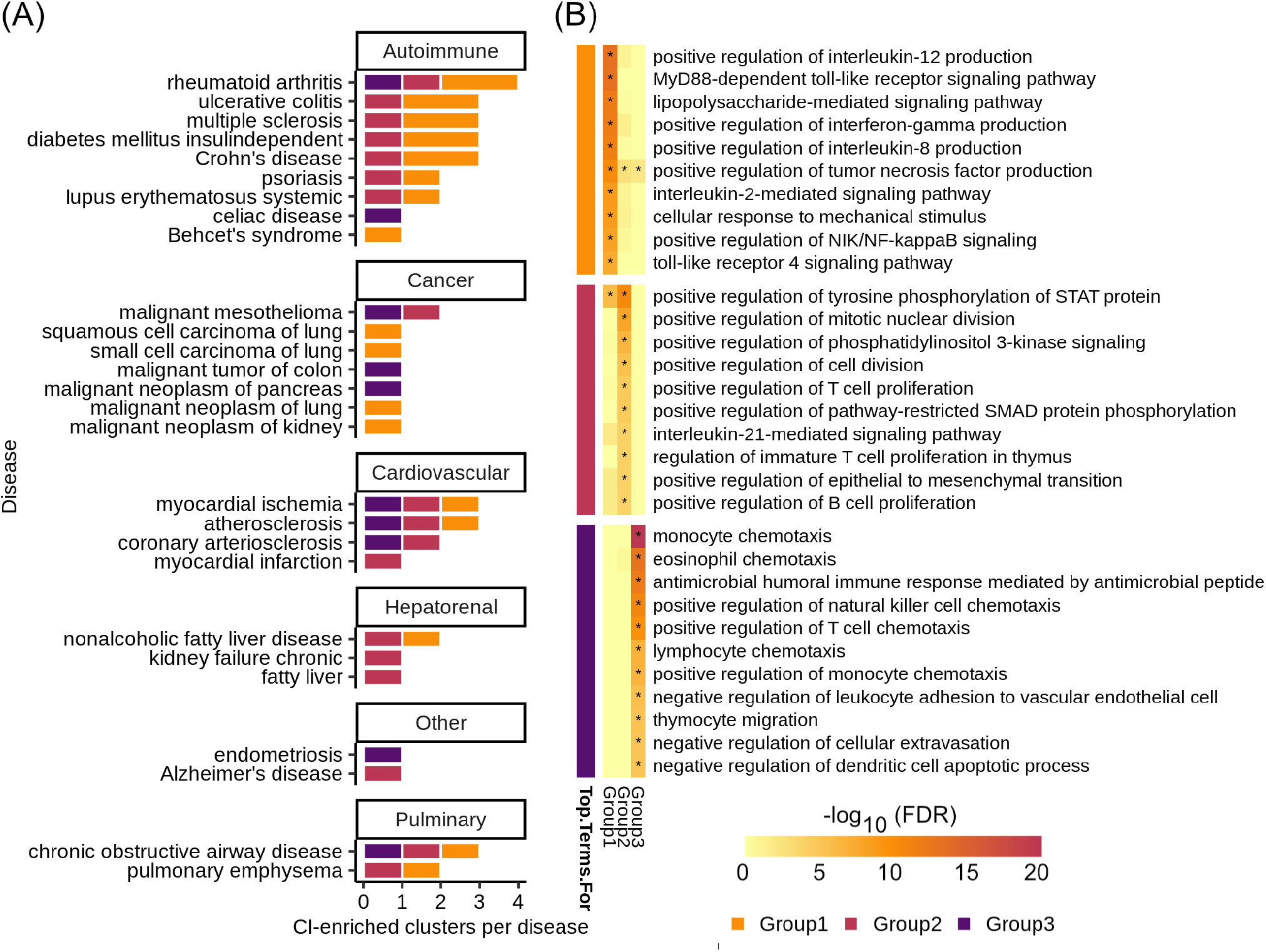
**(A)** Number of CI-enriched clusters per disease colored by CI-signature group. **(B)** Top ten enriched GOBP categories by Benjamini-Hochberg procedure corrected FDR for each CI-signature group — the group is denoted by the colored blocks to the left of the heatmap. The heatmap shows the −*log*_10_(*FDR*) of the enrichment for each CI-signature group — * denotes *FDR* < 0.05.

To determine the biological significance of these signature groups, we performed enrichment analyses for genes unique to each group among GO biological processes (**Fig. 3B, Table S8**). The top 10 significantly enriched terms for each group are largely distinct, with group 1 being enriched for immune relevant signaling pathways, group 2 for regulation of immune cell proliferation, and group 3 for regulation of immune cell chemotaxis (**Fig. 3B**).

### 3.4 Predicting novel treatment opportunities

Our final goal was to leverage the CI-enriched disease clusters we discovered to find potential avenues for repurposing approved drugs to therapeutically target systemic inflammation underlying complex diseases.

Towards this goal, we used SAveRUNNER to find associations between CI-enriched clusters and FDA approved drugs through each drug’s target genes (Fiscon et al. 2021). We found that SAveRUNNER predictions for known treatments were better than random chance —*log*_2_(*auPRC*⁄*prior*) > 0— for disease with more than five known treatments (**Fig. 4A**). Moreover, except for myocardial ischemia, SAveRUNNER predicted drugs in Phase IV clinical trials better than random chance (**Fig. 4A**) (“AACT Database | Clinical Trials Transformation Initiative” n.d.). Drugs in Phase IV are those that have already been proved effective for treating a disease (in Phase III) and are being monitored for long-term safety and efficacy.

**Figure 4:**
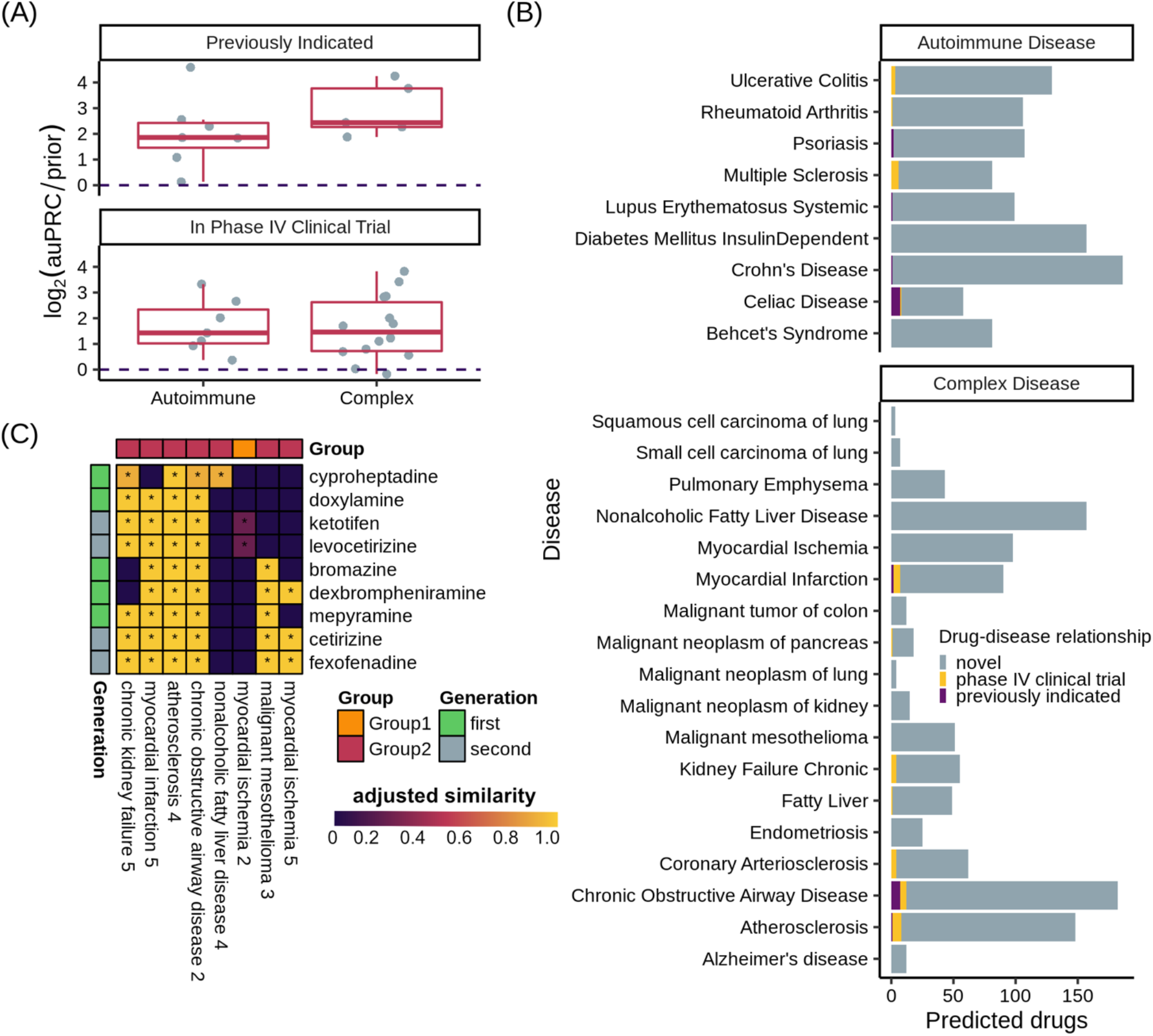
**(A)** log_2_(auPRC/prior) of SAveRUNNER predictions using drugs previously indicated for the disease (top) or drugs ever in Phase IV clinical trials for a disease (bottom) as positive examples. The dotted line is at log_2_(auPRC/prior)= 0. log_2_(auPRC/prior) > 0 denotes predictions better than random chance. **(B)** Number of SAveRUNNER predicted genes (Benjamini-Hochberg procedure corrected *FDR* < 0.01) per disease. **(C)** Heat map showing the adjusted similarity scores between several antihistamines and complex disease clusters — * denotes *FDR* < 0.01. *Group* is the CI-signature the disease clusters is a member of. *Generation* shows whether the drug is a first- or second-generation antihistamine.

SAveRUNNER predicted between 3 and 186 high-confidence (FDR < .01) treatments for each disease and identified previously indicated drugs for 4 of the 9 autoimmune disorders, with significant enrichment among drug predictions for celiac disease (one-sided Fisher’s exact test, BH corrected *FDR* < 0.001, **Fig. 4B, Table S9**.). SAveRUNNER found previously indicated treatments for only 3 of the 18 complex diseases (**Fig. 4B**). This result is expected given that, unlike for autoimmune disorders, most known treatments for these complex disorders are not likely to target the immune system. Treatments previously tested in a clinical trial were predicted for 4 autoimmune disorders and 7 of the complex disorders (**Fig. 4B**).

Many of the top treatment predictions for complex disorders are antihistamines (**Table S6)**, which block the action of histamine at the H^1^-receptor, and the CI-enriched clusters associating antihistamines to diseases mostly reside in the immune-relevant signaling pathway associated CI-signature group 1 (**Fig. 4C**). SAveRUNNER allows for drug prioritization based on the adjusted similarity score between drug target genes and CI-enriched cluster genes. Highly scoring drug-cluster pairs have genes that are closely related in the gene interaction network which increases the likelihood that the drug will be on-target for the paired disease (Fiscon et al. 2021). Multiple antihistamines have equally high adjusted similarity scores for atherosclerosis, suggesting that they have the same potential as treatments for the disease. Conversely, among these top hits, only first-generation antihistamines bromazine and dexbrompheniramine are highly associated with myocardial ischemia, and while cyproheptadine is predicted to treat nonalcoholic fatty liver disease, it is unlikely to be an effective treatment for myocardial infarction or malignant mesothelioma (**Fig. 4C**). This finding suggests that, even among diseases with similar CI-signatures, we can predict disease-specific treatments for the chronic inflammation component of the disease etiology.

## 4 Discussion

Complex diseases exhibit a staggering amount of heterogeneity, being associated with hundreds of genes and with a range of phenotypes. Therefore, to continue advancing our understanding of disease mechanisms and our ability to treat these diseases, it is critical to deconvolve disease heterogeneity by a) resolving subset of disease genes (and cellular processes/pathways) that underlie specific disease-associated phenotypes, and b) identifying avenues to diagnostically and/or therapeutically target those specific phenotypes.

Here, we present a computational data-driven approach to address this critical need (**Fig. 1A**). We used our approach to study chronic inflammation (CI) — a major phenotype in several common, complex diseases. We generated comprehensive lists of (known and predicted) disease-associated genes and identified and classified the CI signal among these genes. We used these signatures to predict novel treatment options to target the inflammatory components of 18 complex diseases.

A key aspect of our approach is ensuring its sensitivity to detect CI disease signatures using autoimmune diseases as positive controls. In autoimmune diseases, the immune system mistakenly attacks healthy tissue causing long-term systemic inflammation. Thus, we expect that the underlying CI disease signatures would be easily identifiable by a valid approach. Indeed, in each of the nine autoimmune diseases analyzed, our approach isolated gene clusters enriched for CI genes (**Fig. 2A**), and identified drugs already used to treat celiac disease, Crohn’s disease, systemic lupus erythematosus, and psoriasis (**Fig. 4B**). This finding is encouraging given that we conservatively matched drugs to diseases only based on expert-curated drug-target data from DGIDb(Freshour et al. 2021) rather than using all drug-target information in DrugCentral(Avram et al. 2021).

To show that our method was not erroneously uncovering CI signals where there were none, we identified 16 UK Biobank traits not patently associated with CI (along with their genes) to use as negative controls. Following this analysis, we found that the median fraction of trait-associated genes predicted by GenePlexus and the median fraction of genes assigned to sizable clusters were lower for these traits than for autoimmune and complex diseases (**Fig. 1D** and **E**). Given that GenePlexus is a method that leverages network connectivity for predicting new genes belonging to a set, these results suggest that the genes associated with non-disease traits may not be as highly connected to one another in ConsensusPathDB as the autoimmune and complex disease genes. Moreover, most of the non-disease trait clusters were not enriched with genes annotated to GO biological processes, suggesting that these clusters are diffuse and that the member genes are unlikely to work together to support a coherent biological task. While non-disease traits like coffee intake and handedness have been associated with inflammation (Paiva et al. 2019; Searleman and Fugagli 1987), this analysis (using GWAS-based trait-associated genes) suggests it is unlikely that SNPs in a coordinated inflammation pathway influence non-disease traits and more likely that any association with inflammation is environmental, not genetic. Taken together, these results suggest that these chosen traits serve as reasonable negative controls and offer a way to meaningfully contrast the results from complex diseases. Ideally, diseases or traits with no underlying inflammatory component but with associated genes that cluster in a network (as well as the autoimmune and complex disease) will serve as better negative controls. Given how common inflammatory processes are in disease, however, such diseases are difficult to definitively identify.

Patients with systemic inflammation caused by one CI-related disorder are more likely to have another CI-related disorder, and, in some cases, treating one can help minimize the risk of diagnosis with a second (Chou et al. 2016; Antohe et al. 2012; Dregan et al. 2014; Armstrong, Harskamp, and Armstrong 2013; Yashiro 2014). To determine how CI-associated disorders relate to one another, we used a network-based approach to quantify the similarity between their CI-enriched clusters. We hypothesized, for example, that Crohn’s disease and “malignant tumor of colon” would have similar CI-signatures, given that patients with inflammatory bowel disease are at increased risk for developing colorectal cancer (S. C. Shah and Itzkowitz 2022). However, Crohn’s disease CI-enriched clusters are members of signature groups 1 and 2, while the “malignant tumor of colon” CI-enriched cluster belongs to group 3 (**Fig. 3A**). Instead of sharing CI-signatures, related CI diseases may, instead, have complementary signatures. Indeed, the group 1 signature, which characterizes two of the three Crohn’s disease CI-enriched clusters, is enriched for genes that positively regulate proinflammatory cytokines like tumor necrosis factor (TNF) and in interferon-gamma (IFNɣ) (**Fig. 3B**). When these cytokines bind to their respective receptors, reactive oxygen species are generated causing oxidative stress (Chatterjee 2016). Oxidative stress, in turn, induces DNA-damage that can induce tumor formation. Colorectal tumors are infiltrated with lymphocytes, which mediate the recruitment of immune cells that suppress tumor growth (Idos et al. 2020). Immune cell infiltration likely leads to our ability to detect the group 3 CI-signature among genes associated with “malignant tumor of colon”, given that group 3 is enriched for immune cell migration and chemotaxis (**Fig. 3A** and **B**). Alternatively, there is a possibility that every CI-associated disease exhibits all three CI-signatures, and our method is only sensitive enough to detect these in a handful of diseases.

We leverage the CI-signatures to identify novel treatment opportunities for the CI-component of 18 complex diseases (**Fig. 4B**). Interestingly, antihistamines were among the top drug associations for 8 of 18 complex diseases (**Fig. 4C**), including atherosclerosis. Atherosclerosis is characterized by the deposition of cholesterol plaques on the inner artery walls. Mast cells, immune cells best known for their response to allergens, are recruited to arteries during plaque progression, where they release histamines. Histamines then activate the histamine H1-receptor, increasing vascular permeability, which allows cholesterol easier access to arteries promoting plaque buildup (Rozenberg et al. 2010). The antihistamines predicted as treatments for CI-related diseases, here, are first and second generation H1-receptor antagonists, which are characterized by the ability to bind peripheral H1-receptors non-selectively (first generation) or selectively (second generation). Mepyramine, one of the first-generation antihistamines highly associated with atherosclerosis (**Fig. 4C**), has already been shown to decrease the formation of atherogenic plaques in a mouse model of atherosclerosis (Rozenberg et al. 2010). Interestingly, it is not predicted as a treatment for myocardial ischemia, which occurs when plaque buildup obstructs blood flow to a coronary artery, suggesting disease-specific antihistamine efficacy even among related diseases. Second generation antihistamines cetirizine and fexofenadine are also highly associated with atherosclerosis but neither prevented or reduced atherosclerosis progression in a mouse model of atherosclerosis, and both increased atherosclertotic lesions at low doses (Raveendran et al. 2014). In the expert curated drug target database used in this study, the H1-receptor is the only target listed for all three drugs suggesting that drug-specific off-target effects are mediating atherosclerosis treatment outcomes. Nevertheless, we generated a list of many other high priority drugs that may be effective in treating the CI-component of atherosclerosis and other complex diseases.

Overall, we have shown that our method can isolate the chronic inflammation gene signature of a complex disease using a network-based strategy and, by integrating information across multiple complementary sources of data, it can predict and prioritize potential therapies for the systemic inflammation involved in that disease specifically. Importantly, our approach provides a blueprint for identifying and prioritizing therapeutic opportunities for any disease endophenotype.

## 5 Contribution to the field

Complex disorders such as cardiovascular diseases, diabetes, cancer, and Alzheimer’s disease exhibit a staggering amount of heterogeneity, each disorder being associated with hundreds of genes and with a range of phenotypes. One such phenotype associated with many of these disorders is systemic chronic inflammation. Chronic inflammation occurs when our body’s inflammatory responses do not resolve, resulting in persistent, low-grade immune activation that causes collateral damage to the affected tissue over time. To advance our understanding of this common phenotype and our ability to treat the resulting damage in each disease, it is critical to find subsets of disease genes that underlie chronic inflammation and then identify avenues to diagnostically and/or therapeutically target those genes. Here, we present a novel computational approach to achieve this goal using a molecular network that captures how pairs of genes in the human genome interact with each other to perform their functions inside cells in our body. Our approach, built based on machine learning and clustering, uses the molecular network and multiple other sources of data to isolate gene “signatures” of chronic inflammation specific to each disorder. We quantify the similarity between inflammation signatures across diseases and predict and prioritize potential new treatment opportunities specifically targeting the ‘inflammation’ component of complex diseases. Importantly, our strategy can be used to pinpoint the gene signature of any phenotype underlying a complex disease and identify drugs to treat that specific phenotype.

## 7 Author contributions

SLH, AM, and AK conceived and designed the approach and experiments and wrote the manuscript; SLH and AM implemented the approach and performed the experiments. CAM wrote the GenePlexus code used in the experiments. All authors read, edited, and approved the final manuscript.

## 9 Acknowledgements

We thank the members of the Krishnan Lab for helpful discussions.

## 11 Data availability statement

The data, main and supplemental results, and code used to reproduce this study are freely available at https://github.com/krishnanlab/chronic-inflammation.

