## Supplementary Data, Figures, and Table legends for "A network-based approach for isolating the chronic inflammation gene signatures underlying complex diseases towards finding new treatment opportunities"

#### Supplementary Material

#### 1 Supplementary Data

The data, main and supplemental results, and code used to reproduce this study are freely available at https://github.com/krishnanlab/chronic-inflammation.

#### 2 Supplementary Tables

##### 2.1 Table S1: Disease and trait GenePlexus average $log_2\left(\frac{auPRC}{prior}\right)$ from 3-fold cross validation – all networks

Column Descriptions:

1. Disease: Disease or non-disease trait
2. PredictionNetwork: Network that GenePlexus used to make predictions
3. Features: GenePlexus used the columns of the network's adjacency matrix as features in each case.
4. negativesFrom: Denotes if the negative example genes were derived from the Gene Ontology (GO) or from DisGeNet.
5. nSeeds: Number of genes originally associated with the disease/trait by DisGeNet/Pascal
6. nNegs: Number of negative example genes
7. Threshold: The cut off threshold for genes newly associated to the disease/trait by GenePlexus (> 0.80 in all cases)
8. CVscore: Average $log_2\left(\frac{auPRC}{prior}\right)$ from 3-fold cross validation
9. TraitType: Denotes if the disease/trait is a "Non-disease Trait", "Complex Disease", "Autoimmune Disease", or "Inflammation Gene Set".

##### 2.2 Table S2: Disease and trait seed genes

Genes originally associated with a disease by DisGeNet or a non-disease trait by Pascal

Column Descriptions:

1. GeneSymbol: The gene symbol
2. EntrezID: The gene Entrez ID

3. Disease: Disease or non-disease trait
4. TraitType: Denotes if the disease/trait is a “Non-disease Trait”, “Complex Disease”, or “Autoimmune Disease”.

#### 2.3 Table S3: Gene/Cluster assignments – all network/CI gene set combinations

Column Descriptions:

1. PredictionNetwork: Network that GenePlexus used to make predictions
2. ClusterGraph: Network used to make and cluster disease-specific subgraphs
3. Disease: Disease or non-disease trait
4. TraitType: Denotes if the disease/trait is a “Non-disease Trait”, “Complex Disease”, or “Autoimmune Disease”.
5. nClusteredPermutations: The number of randomly permuted gene lists (out of 5,000) for each disease that clustered with 100 iterations of the Leiden algorithm. If no randomly permuted gene lists clustered the disease/trait was not included in the subsequent analyses.
6. Cluster: Cluster assignment
7. Entrez: The gene Entrez ID
8. Symbol: The gene symbol
9. Probability: The GenePlexus predicted probability that the gene is associated with the disease/trait
10. GeneType: Denotes if the gene is predicted by GenePlexus to be associated with the disease/trait or an original seed gene.

#### 2.4 Table S4: CI gene sets with GenePlexus predictions

We used GenePlexus to predict the association of every gene in the network of interest with a supplied chronic inflammation gene set. Genes with a probability $\geq$ 0.80 were included in the expanded gene list. Related to Figures 2C and S3-7.

Column Descriptions:

1. Entrez: The gene Entrez ID
2. Symbol: The gene symbol
3. Probability: The GenePlexus predicted probability that the gene is associated with the chronic inflammation gene set
4. ChronicInflammationSource: The source of the chronic inflammation gene set
5. PredictionNetwork: Network that GenePlexus used to make predictions
6. GeneType: Denotes if the gene is an original seed gene, predicted by GenePlexus to be associated with the chronic inflammation gene set (probability $\geq$ 0.80), or not likely to be CI-associated (probability $<$ 0.80).

#### 2.5 Table S5: CI-overlap results – all network/CI gene set combinations

Column Descriptions:

1. PredictionNetwork: Network that GenePlexus used to make predictions
2. ClusterGraph: Network used to make and cluster disease-specific subgraphs

3. ChronicInflammationSource: The source of the chronic inflammation gene set
4. Disease: Disease or non-disease trait
5. Cluster: Cluster assignment
6. nOverlap: Intersection of genes in the cluster and the CI gene set
7. nChronicInflammationSourceGenes: Number of genes in the CI gene set
8. nClusterGenes: Number of genes in the cluster
9. Enrichment: Enrichment score (see *Methods*)
10. PermutedPval: Permutation test-based p-value (see *Methods*)
11. PermutedFDR: Within disease BH-corrected p-value
12. nClustersFromPermutedSets: Number of permuted clusters used to calculate PermutedPval

#### 2.6 Table S6: Number of diseases considered for CI enrichment analysis – all network/CI gene set combinations

Diseases/traits were only included in the CI enrichment analysis if:

- The GenePlexus average $log_2\left(\frac{auPRC}{prior}\right) \geq 1$ (CV score)
- At least one randomly permuted gene list associated with the disease had clusters containing $\geq 5$ genes.
- At least one real cluster contained $\geq 5$ genes

Column Descriptions:

1. Trait Type: Denotes if the disease/trait is a "Non-disease Trait", "Complex Disease" or "Autoimmune Disease"
2. Prediction Network: Network that GenePlexus used to make predictions
3. Total: The total number of original diseases/traits considered for analysis
4. CV score ≥ 1: Number of traits with a GenePlexus model with averag $\mathrm{log2(auPRC/prior)} \geq 1$
5. Cluster Graph: Network used to make and cluster disease-specific subgraphs
6. Permuted Clusters: Number of diseases/traits with at least one randomly permuted gene list associated with the disease had clusters containing $\geq 5$ genes
7. Clusters: Number of disease/traits with at least one real cluster containing $\geq 5$ genes
8. Final: Final number of disease/traits used in the CI enrichment analysis

#### 2.7 Table S7: CI-signature group assignments – ConsensusPathDB/Geneshot with more than 10 publications only

The CI-enriched clusters from every disease were grouped based on their pair-wise adjusted similarity scores calculated using the SAveRUNNER algorithm. The cluster to CI-signature group assignments are listed here. Related to Fig 3.

Column Descriptions:

1. Cluster: CI-enriched cluster
2. CI_SignatureGroup: Group assignment
3. Disease: Disease subnetwork the cluster is derived from
4. TraitType: Denotes if the disease is a "Complex Disease" or "Autoimmune Disease".

5. TraitTypeFine: Denotes which finer disease type a disease belongs to: “Cardiovascular”, “Autoimmune Disease”, “Pulminary”, “Hepatorenal”, “Cancer”, or “Other”.

#### 2.8 Table S8: GOBP enrichments for CI-signature groups – ConsensusPathDB/Geneshot with more than 10 publications only

Column Descriptions:

1. CI_SignatureGroup: group the gene set is derived from
2. GO.ID: The ID associated with the GOBP term
3. Term: GOBP term name
4. Annotated: Total number of genes annotated to that term
5. Significant: Number of genes in the gene set intersecting the genes annotated to the term
6. Expected: Expected proportion of genes in a random gene set intersecting the genes annotated to the term
7. Pval: Fisher test p-value
8. FDR: Within signature group BH-corrected p-values
9. OverlappingGenes: Entrez IDs for the intersecting genes
10. Log10FDR: $-log_{10}(FDR)$

#### 2.9 Table S9: SAveRUNNER results – ConsensusPathDB/Geneshot with more than 10 publications only

Column Descriptions:

1. Disease: Disease associated with the drug
2. TraitType: Denotes if the disease is a “Complex Disease” or “Autoimmune Disease”.
3. Cluster: CI-enriched cluster through which the drug is associated
4. Drug: Associated drug
5. Pval: SAveRUNNER p-value
6. FDR: BH-corrected p-value
7. Known_Relationship: Denotes if a disease’s relationship to a drug is “previously indicated”, “off-label use”, or “none”
8. PairInCT: TRUE if the drug-disease pair has been tested in a Phase IV clinical trial
9. Adjusted_similarity: Similarity score between the drug and disease calculated by SAveRUNNER

### 3 Supplementary Figures

#### 3.1 Figure S1: Number of genes in each inflammation gene set

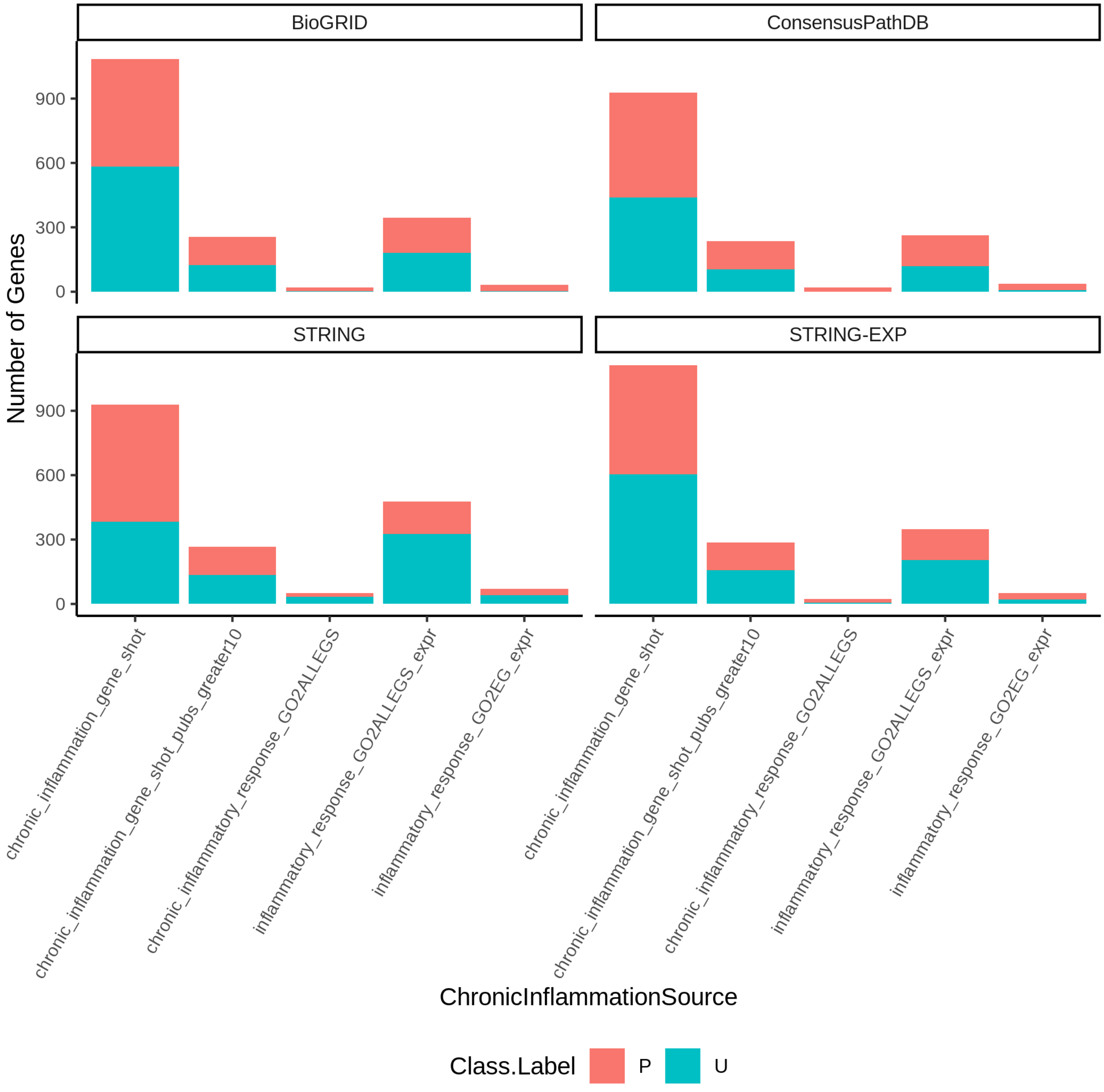


**Figure S1.** Number of CI-associated genes per CI gene set source, predicted on each network. "P" denotes positive example genes for GenePlexus. These are the original seed genes. "U" denotes genes predicted by GenePlexus to be associated with the CI gene set with a probability ≥ 0.80

#### 3.2 Figure S2: Proportion of traits overlapping with at least one chronic inflammation cluster – all network/CI gene set combinations

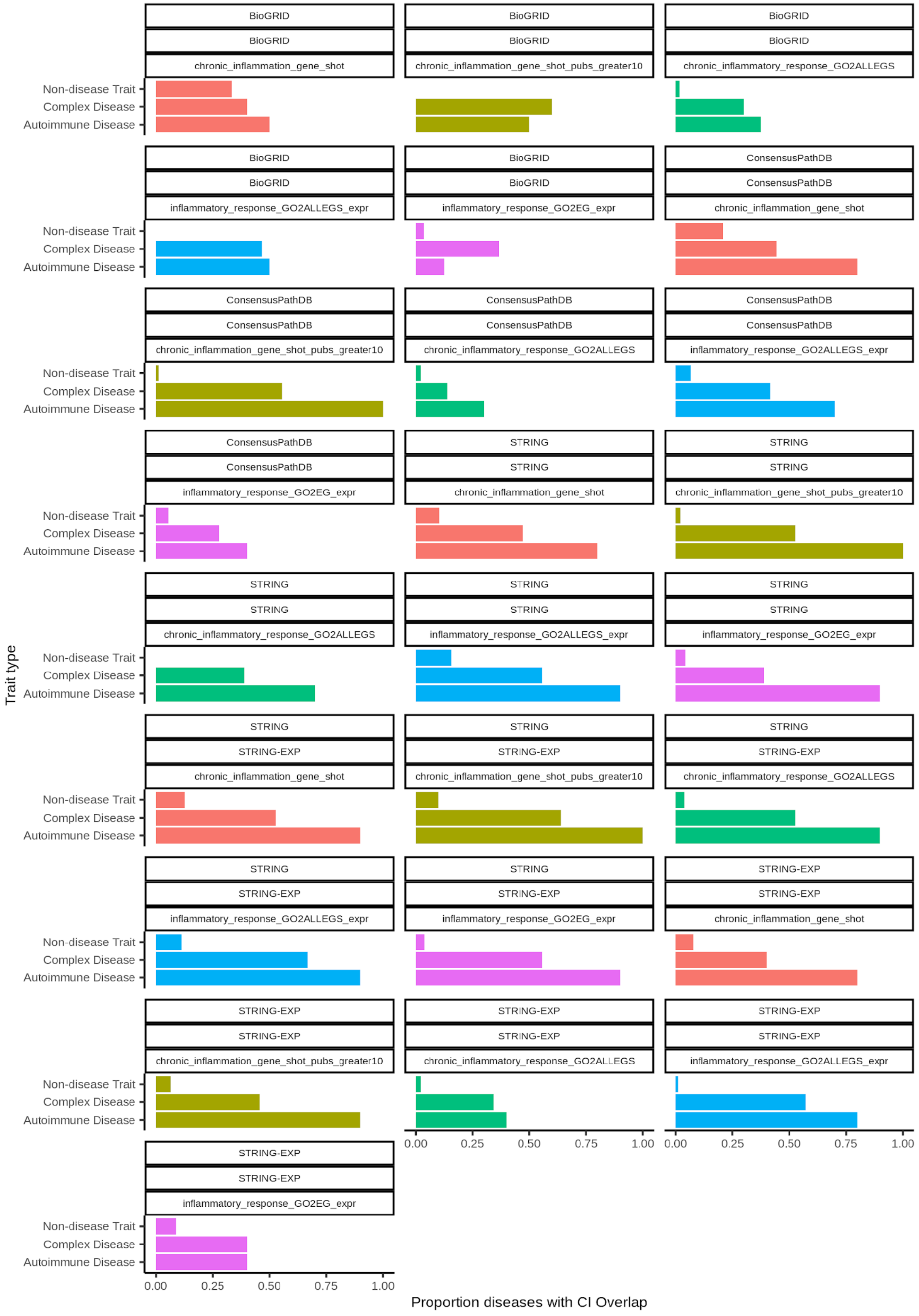

**Figure S2.** Proportion of diseases/traits with at least one significant CI-overlapping cluster. Above each plot is top) the prediction network used by GenePlexus, middle) the network used for clustering, and bottom) the CI gene set source.

#### 3.3 Figure S3: Chronic inflammation association score of non-seed genes in overlapping and non-overlapping clusters – predicted and clustered on BioGRID

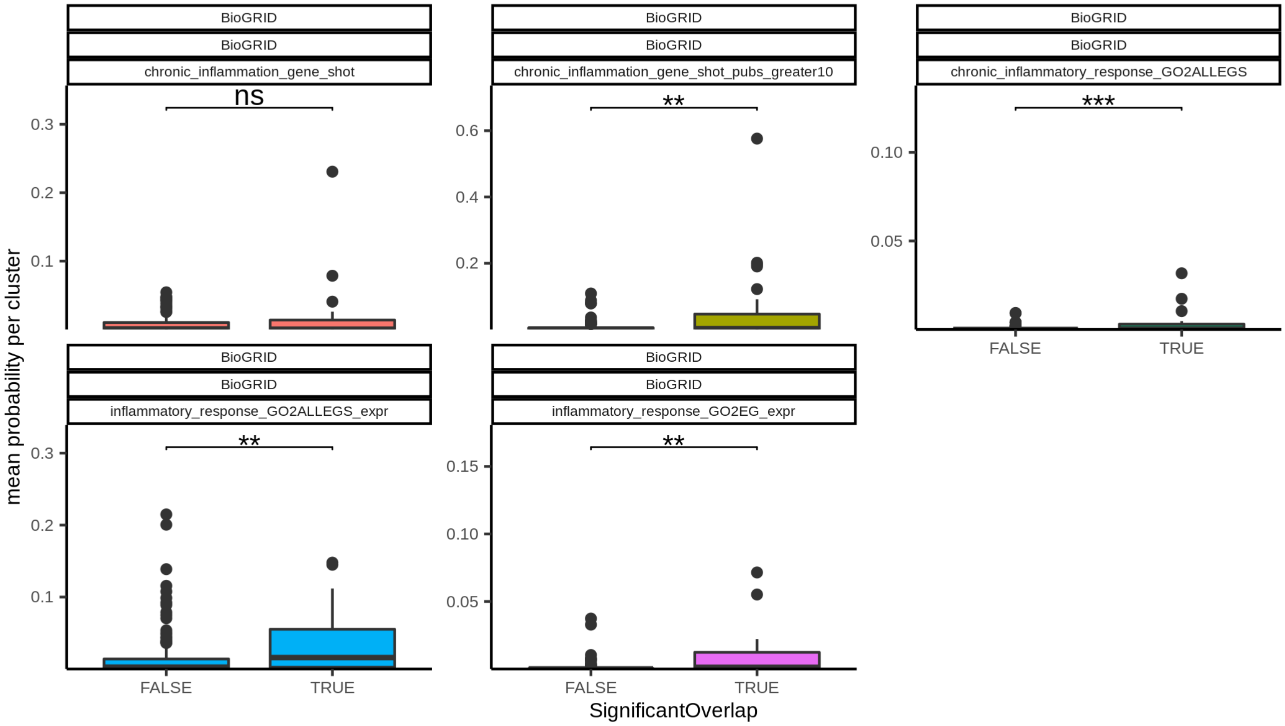


**Figure S3.** Mean probability that genes with no known relationship with chronic inflammation residing in a CI-enriched cluster or non-CI-enriched cluster are associated with CI. FDR was calculated using a BH-adjusted Wilcoxon signed-rank test. (ns $p >= 0.05$, $* \; p < 0.05$, $** \; p < 0.01$, $*** \; p < 0.001$, $**** p < 1x10^{-4}$). Above each plot is top) the prediction network used by GenePlexus, middle) the network used for clustering, and bottom) the CI gene set source.

##### 3.4 Figure S4: Chronic inflammation association score of non-seed genes in overlapping and non-overlapping clusters – predicted and clustered on ConsensusPathDB

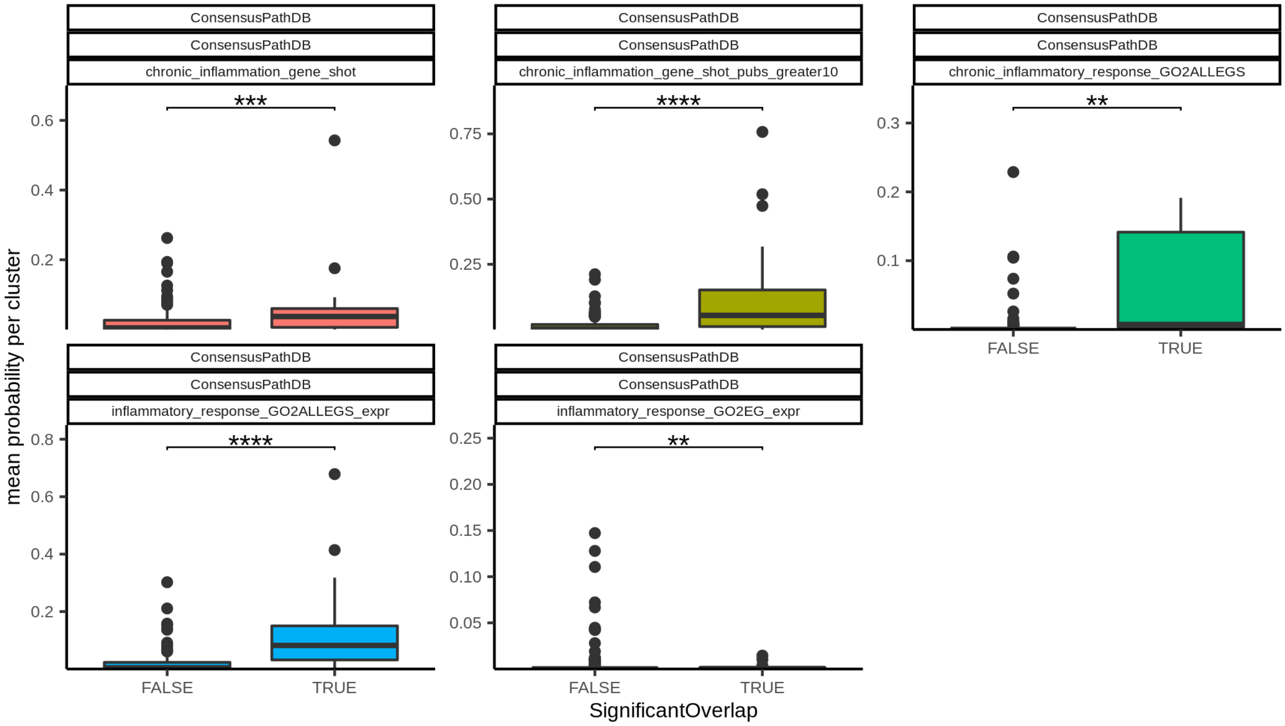


**Figure S4.** Mean probability that genes with no known relationship with chronic inflammation residing in a CI-enriched cluster or non-CI-enriched cluster are associated with CI. FDR was calculated using a BH-adjusted Wilcoxon signed-rank test. ($* \, p < 0.05$, $** \, p < 0.01$, $*** \, p < 0.001$, $**** p < 1x10^{-4}$). Above each plot is top) the prediction network used by GenePlexus, middle) the network used for clustering, and bottom) the CI gene set source.

#### 3.5 Figure S5: Chronic inflammation association score of non-seed genes in overlapping and non-overlapping clusters – predicted and clustered on STRING-EXP

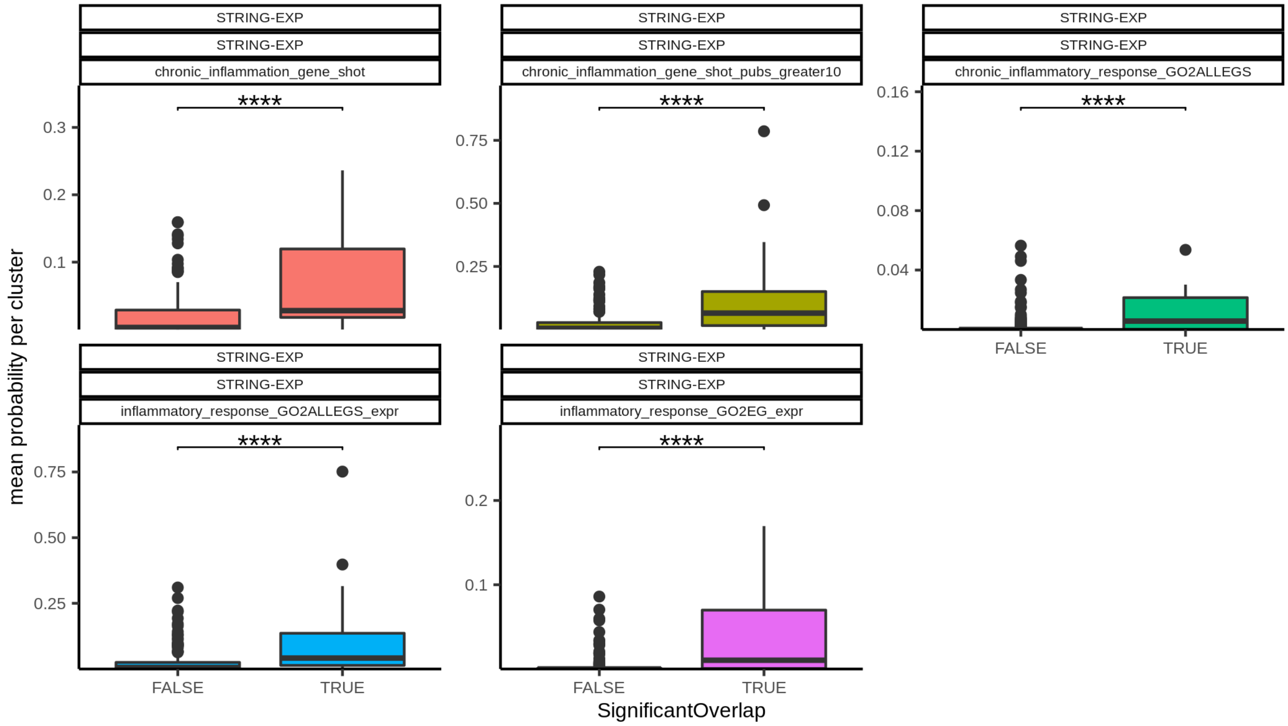


**Figure S5.** Mean probability that genes with no known relationship with chronic inflammation residing in a CI-enriched cluster or non-CI-enriched cluster are associated with CI. FDR was calculated using a BH-adjusted Wilcoxon signed-rank test. ($* \, p < 0.05$, $** \, p < 0.01$, $*** \, p < 0.001$, $**** p < 1x10^{-4}$). Above each plot is top) the prediction network used by GenePlexus, middle) the network used for clustering, and bottom) the CI gene set source.

#### 3.6 Figure S6: Chronic inflammation association score of non-seed genes in overlapping and non-overlapping clusters – predicted and clustered on STRING

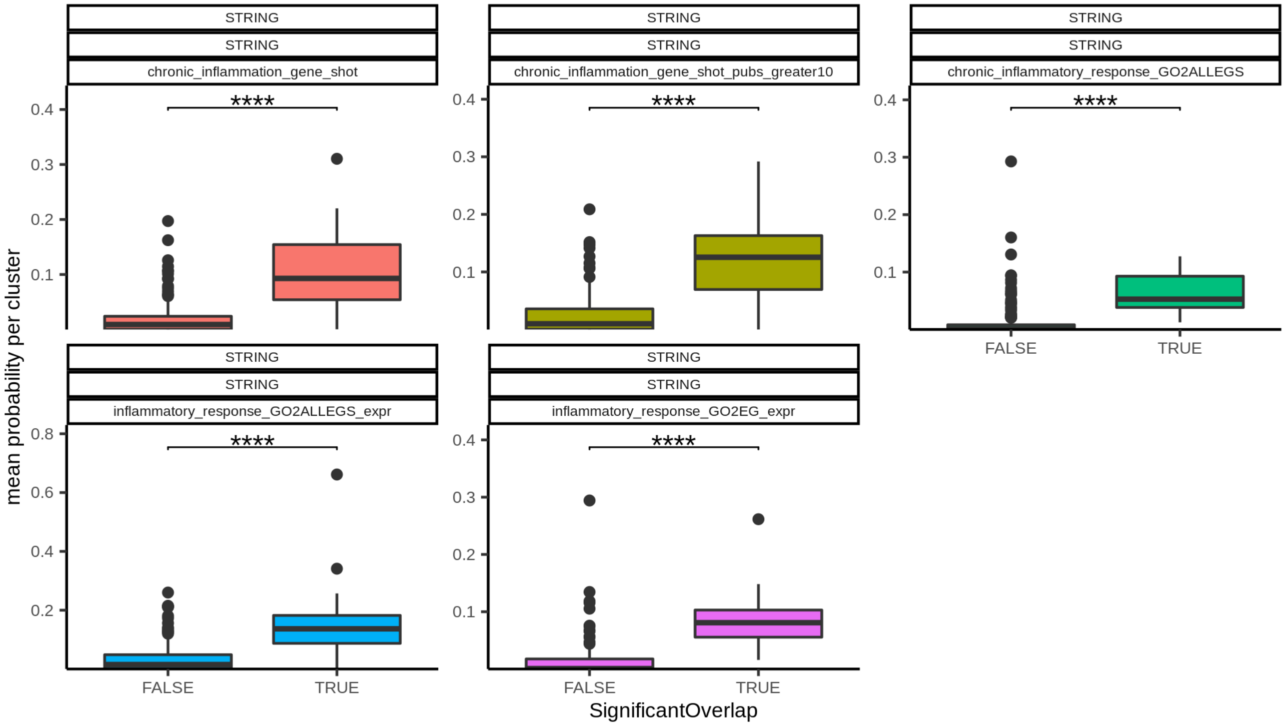


**Figure S6.** Mean probability that genes with no known relationship with chronic inflammation residing in a CI-enriched cluster or non-CI-enriched cluster are associated with CI. FDR was calculated using a BH-adjusted Wilcoxon signed-rank test. ($* \, p < 0.05$, $** \, p < 0.01$, $*** \, p < 0.001$, $****p < 1x10^{-4}$). Above each plot is top) the prediction network used by GenePlexus, middle) the network used for clustering, and bottom) the CI gene set source.

#### 3.7 Figure S7: Chronic inflammation association score of non-seed genes in overlapping and non-overlapping clusters – predicted on STRING and clustered on STRING-EXP

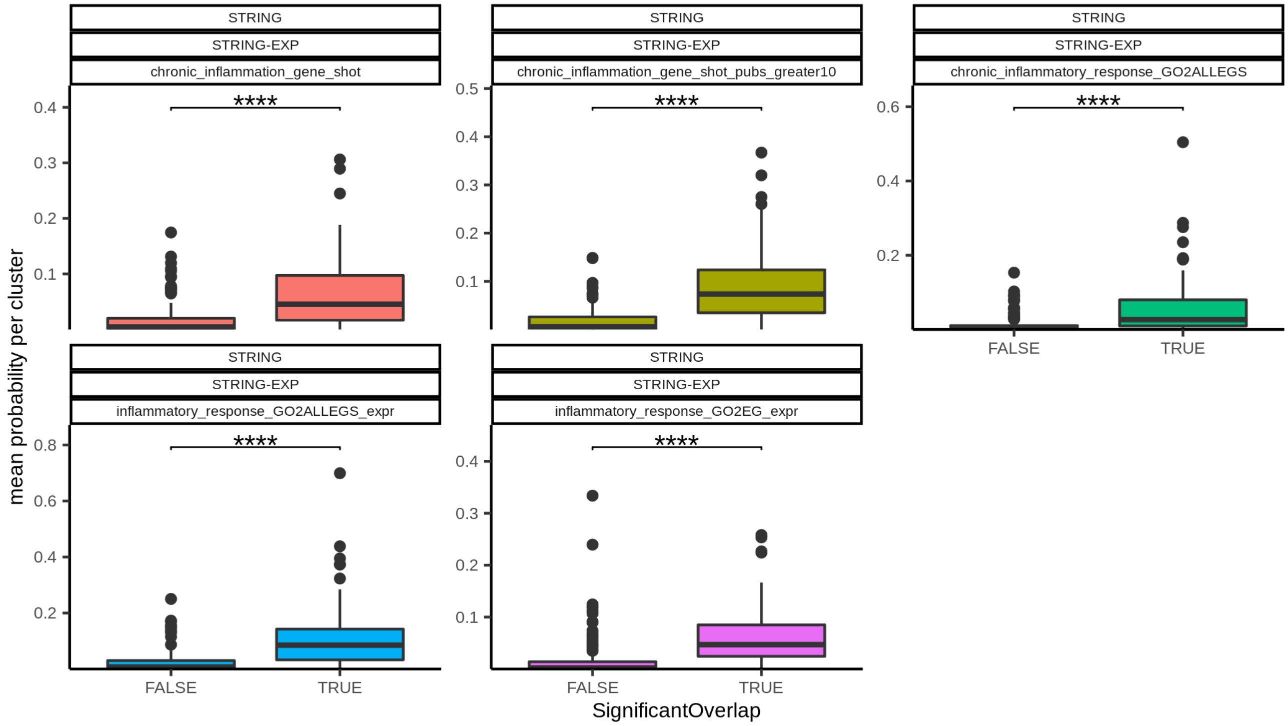


**Figure S7.** Mean probability that genes with no known relationship with chronic inflammation residing in a CI-enriched cluster or non-CI-enriched cluster are associated with CI. FDR was calculated using a BH-adjusted Wilcoxon signed-rank test. ($*\ p < 0.05$, $**\ p < 0.01$, $***\ p < 0.001$, $****p < 1x10^{-4}$). Above each plot is top) the prediction network used by GenePlexus, middle) the network used for clustering, and bottom) the CI gene set source.
